# Nuclear Factor Kappa B Over-Activation in the Intervertebral Disc Leads to Macrophage Recruitment and Severe Disc Degeneration

**DOI:** 10.1101/2023.08.07.552274

**Authors:** Kevin G. Burt, Min Kyu M. Kim, Dan C. Viola, Adam C. Abraham, Nadeen O. Chahine

## Abstract

**Objective:** Low back pain (LBP) is the leading cause of global disability and is thought to be driven primarily by intervertebral disc (IVD) degeneration (DD). Persistent upregulation of catabolic enzymes and inflammatory mediators have been associated with severe cases of DD. Nuclear factor kappa B (NF-κB) is a master transcription regulator of immune responses and is over expressed during inflammatory-driven musculoskeletal diseases, including DD. However, its role in triggering DD is unknown. Therefore, this study investigated the effect of NF-κB pathway over-activation on IVD integrity and DD pathology.

**Methods:** Using skeletally mature mouse model, we genetically targeted IVD cells for canonical NF-κB pathway activation via expression of a constitutively active form of inhibitor of κB kinase B (IKKβ), and assessed changes in IVD cellularity, structural integrity including histology, disc height, and extracellular matrix (ECM) biochemistry, biomechanics, expression of inflammatory, catabolic, and neurotropic mediators, and changes in macrophage subsets, longitudinally up to 6-months post activation.

**Results:** Prolonged NF-κB activation led to severe structural degeneration, with a loss of glycosaminoglycan (GAG) content and complete loss of nucleus pulposus (NP) cellularity. Structural and compositional changes decreased IVD height and compressive mechanical properties with prolonged NF-κB activation. These alterations were accompanied by increases in gene expression of inflammatory molecules (*Il1b, Il6, Nos2*), chemokines (*Mcp1*, *Mif*), catabolic enzymes (*Mmp3, Mmp9, Adamts4*), and neurotrophic factors (*Bdnf*, *Ngf*) within IVD tissue. Increased recruitment of activated *F4/80*^+^ macrophages exhibited a greater abundance of pro-inflammatory (CD38^+^) over inflammatory-resolving (CD206^+^) macrophage subsets in the IVD, with temporal changes in the relative abundance of macrophage subsets over time, providing evidence for temporal regulation of macrophage polarization in DD *in vivo,* where macrophages participate in resolving the inflammatory cascade but promote fibrotic transformation of the IVD matrix. We further show that NF-κB driven secretory factors from IVD cells increase macrophage migration and inflammatory activation, and that the secretome of inflammatory-resolving macrophages mitigates effects of NF-κB overactivation.

**Conclusion:** Overall the observed results suggest prolonged NF-κB activation can induce severe DD, acting through increases in inflammatory cytokines, chemotactic proteins, catabolic enzymes, and the recruitment and inflammatory activation of a macrophage cell populations, that can be mitigated with inflammatory-resolving macrophage secretome.

## INTRODUCTION

Chronic inflammation plays a critical role across musculoskeletal tissue diseases by contributing to degeneration and pain, which have a massive global impact on disability and wellbeing. Among musculoskeletal tissue diseases, low back pain (LBP) is the leading cause of disability globally, where a growing prevalence and limited therapeutic interventions drives an annual U.S. economic encumbrance over $100 billion (1, 2). Although the etiology of LBP is multifactorial, intervertebral disc (IVD) degeneration is the most prevalent contributor to symptomatic LBP (3). A commonly proposed signaling driver of chronic musculoskeletal inflammation is the master transcription factor nuclear factor kappa B (NF-κB), which is known to mediate immune responses and has been observed to be elevated locally in connective tissues of patients with disc degeneration (DD) (4), and other musculoskeletal soft tissue diseases such as tendinopathy (5), knee osteoarthritis (OA) (6), and synovial rheumatoid arthritis (RA) (7).

Molecularly, canonical NF-κB activation is mediated by degradation of the inhibitor of NF-κB (IκBα) by the IκB kinase (IKK) complex, containing the regulating subunit, IKKγ, and catalytic subunits, IKKα and IKKβ. IκBα degradation allows NF-κB to freely translocate from the cytoplasm to the nucleus to regulate transcription. During canonical pathway activation NF-κB subunits, most commonly p50 and p65, regulate transcription of downstream inflammatory signaling (8, 9). Within the IVD and during degeneration, inflammatory cytokines (10, 11) and catabolic enzymes (12–14) have been found to be persistently elevated in human degenerated IVDs. Coinciding with persistent inflammation, increased innate immune cell presence, specifically macrophages, has been observed in DD samples and is thought to engage in inflammatory driven crosstalk with IVD cells (15–17). The master transcription factor, NF-κB, is known to regulate a number of these inflammatory cytokines and catabolic enzymes, as well as mediate immune cell recruitment and activation (8, 18, 19).

Though chronic inflammation is thought to be a key driver of DD, there is a lack of *in vivo* models of inflammatory driven DD. Surprisingly, prior studies evaluating global inflammatory mutations have yielded mixed outcomes on the effects to IVD integrity (20, 21). Specifically, investigations of global overexpression of human *TNFα* in Tg197 mice resulted in systemic inflammation with higher incidence of spontaneous herniation, increased immune cell presence, and degenerative changes in vertebral bone, but no overt evidence of DD was observed, suggesting that human *TNFα* driven systemic inflammation does not produce severe DD (20). In a model of IL-1 mediated inflammation, IL-1 receptor antagonist (IL-1ra) knockout mice were found to exhibit features of DD (structural degeneration, increased catabolic enzyme expression), suggesting that deletion of the natural inhibitor of IL-1 (IL-1ra) created a global IL-1 imbalance, which could serve as a possible driver of DD (21). Yet, another study that investigated IVD integrity in IL-1α/β knock out mice found little to no effect on IVD integrity, despite evidence of decreased systemic cytokine levels (22). Moreover, global deletion of IL-1ra had no appreciable effect on the response of IVD to puncture injury (23). Together the unexpected and disparate effects observed using global over activation or knock out models highlight the complexity of using regulation of systemic inflammatory signaling to study local effects on IVD integrity, in part because the IVD naturally exists in an avascular niche, which may limit the impact of systemic inflammation on the IVD. Therefore, there remains a gap in knowledge on whether persistent local IVD inflammation can produce severe DD *in vivo*.

To directly investigate the role of discal inflammation in the development of DD we utilized an inducible cartilage specific genetic mouse model to target constitutive activation of NF-κB in all compartments of the IVD, assessing for tissue structural, compositional, and biological changes. We hypothesized that prolonged canonical NF-κB pathway activation within IVD cells will induce a chronic pro-inflammatory microenvironment that mimics inflammatory features of clinical DD, which in turn will activate an innate immune response and produce advanced morphological DD in otherwise healthy adult mice. Findings indicate that inducing persistent inflammation in the IVD is sufficient to cause severe DD mediated by increased pro-inflammatory cytokine, chemokine, and catabolic enzyme expression and macrophage recruitment. Furthermore, we demonstrate the secretome of IVD cells over-expressing IKKβ directly promote an inflammatory macrophage phenotype, and that this modulation could be mitigated by paracrine factors derived from inflammatory-resolving macrophages.

## RESULTS

### Development and validation of sustained IKKβ-NF-κB over-activation within the IVD

We used a mouse carrying the inducible Cre recombinase construct, CreERT2, in the aggrecan (*Acan*) gene to target signaling specifically in IVD *Acan^+^* cells. To validate the presence of Cre-mediated genetic recombination within the adult IVD, a *Acan^CreERT2^* mouse was crossed with a *Ai14* fluorescent reporter mouse (*AcanCre;Ai14*) and IVDs were evaluated following IP tamoxifen injections. Red fluorescent protein (RFP) expression indicative of Cre-activity was detected throughout all tissue compartments of the caudal IVD (nucleus pulposus (NP), annulus fibrosis (AF), cartilage endplate (EP)) and in the vertebral growth plates (GP) at 3-days (Fig. 1B) and 3-months (Fig. 1D) following IP tamoxifen injections (or post recombination). No RFP expression was observed in the *AcanCre*^-/-^ mice (Fig. 1A,C). Following successful validation that *Acan^CreERT2^*mice target IVD tissues for genetic recombination, we next crossed *Acan^CreERT2^* mice with *Ikk2ca^fl/fl^*mice to induce IKKβ-NF-κB over-activation in *Acan*^+^ IVD cells upon tamoxifen IP injection (*AcanCre^ERT2/+^;Ikk2ca^fl/fl^*, referred to from here on as IKKβCA mice). IKKβ expression was assessed in IKKβCA mice compared to the control mice (*Acan^+/+^;Ikk2ca^fl/fl^*) at 1-month post recombination. Staining for cells positive for IKKβ were detected in all IVD tissue types (NP, AF and EP) from IKKβCA mice (Fig. 2A), in a pattern similar to the Cre activity observed in reporter mice. No detectable staining for IKKβ was seen in control IVDs (Fig. 2A). Moreover, significantly increased *Ikk2* gene expression was detected in the IVDs of IKKβCA mice shortly following recombination, at 1-week (p=0.0014), with sustained increase in *Ikk2* gene expression at 2-months (p=0.0014) post recombination (Fig. 2B). Similar results in Cre-activity and increases in IKKβ protein and gene (*Ikk2*) expression were also observed within lumbar IVDs (Fig. S1). Validation results showed this *in vivo* model effectively targets all IVD tissue compartments producing an increased and sustained IKKβ expression at both gene and protein levels.

**Figure 1:**
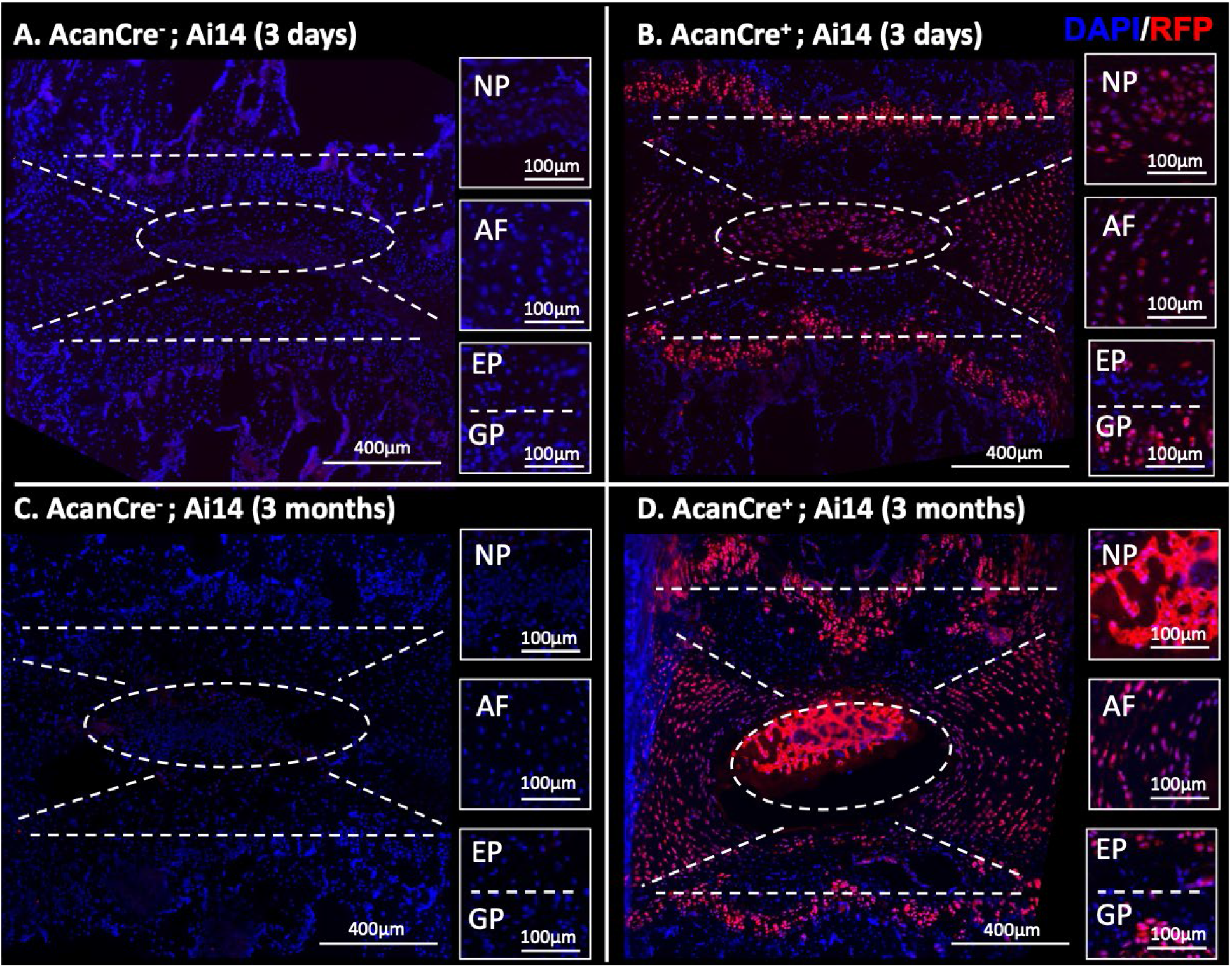
The *Acan^CreERT2^* mouse targets all compartments of the IVD and GP. Representative images of IF staining for RFP within mid sagittal sections of (**A,C**) *AcanCre^-^;Ai14* and (**B,D**) *AcanCre^+^;Ai14* reporter mice 3-days and 3-months following tamoxifen IP injections. NP, AF, EP, GP compartments are delineated (white dashed lines).

**Figure 2:**
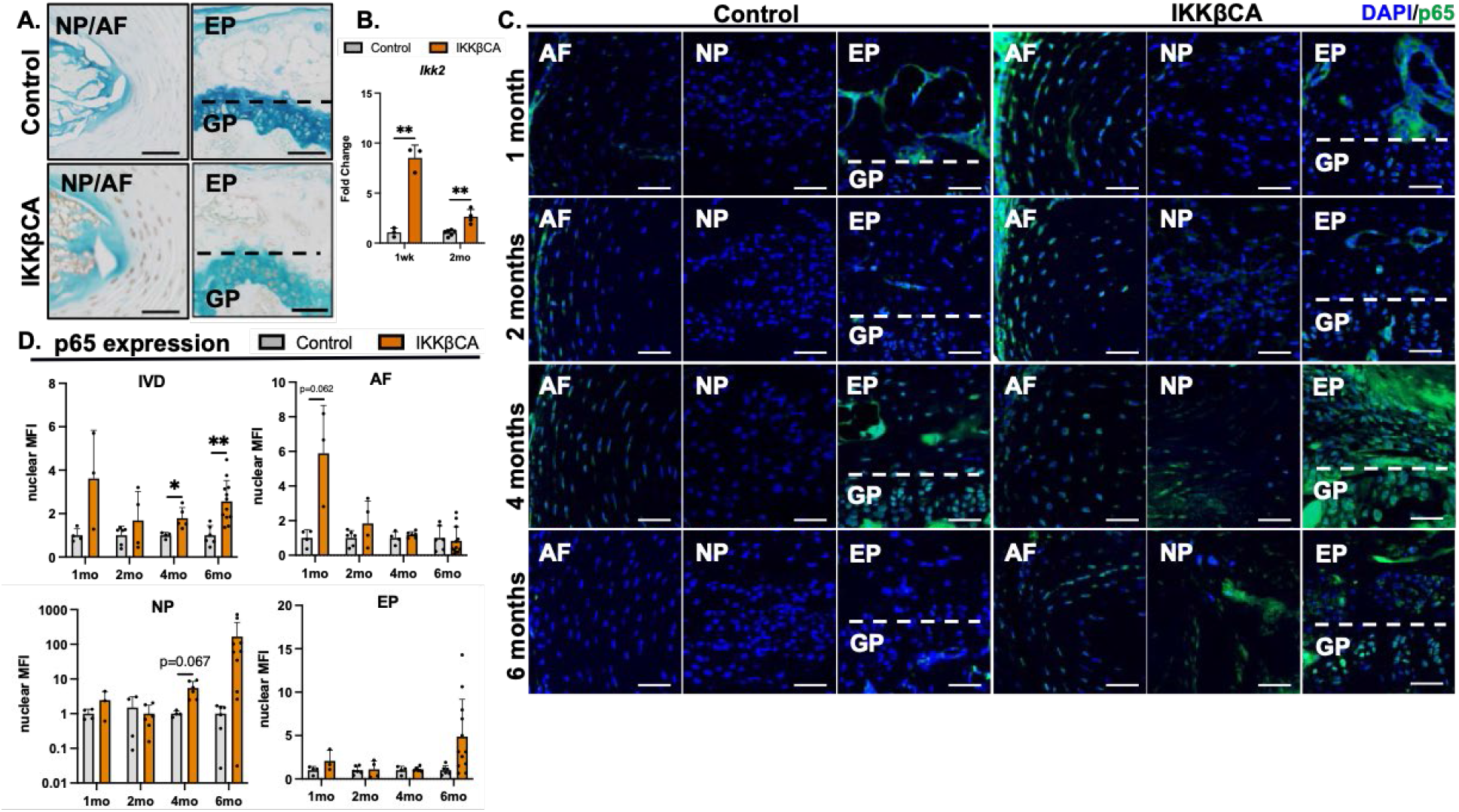
IKKβ over-expression and NF-κB activation within IKKβCA mice. (**A**) Representative images of IHC staining for IKKβ within mid sagittal sections of 1-month control and IKKβCA discs. Scale bar = 100μm. (**B**) Gene expression of *Ikk2* within IVDs expressed as fold change relative to control. (**C**) Representative IF staining for phosphorylated p65 (green) within mid sagittal sections of control and IKKβCA discs. Scale bar = 50μm. (**D**) Nuclear MFI quantification (normalized to control within time point) of phosphorylated p65. *p<0.05, **p<0.01.

We then assessed activation of p65 subunit in the canonical IKKβ-NF-κB pathway, with staining for phosphorylated p65 (phospho-p65) (8). Increased immunofluorescence staining of phospho-p65 was detected in all three IVD tissues of IKKβCA mice compared to control mice at 4-(p=0.048) and 6-months (p=0.0073) post activation (Fig. 2C,D). Analysis of p65 activation within tissue compartments revealed a trending increases in phospho-p65 nuclear MFI in the AF at 1-month (p=0.062) and in the NP at 4-months (p=0.067) post recombination in IKKβCA mice compared to control mice (Fig. 2C,D). These results demonstrate that this model produces expression of constitutively active IKKβ within the IVD, resulting in increased and prolonged activation of downstream canonical NF-κB signaling pathway activation.

### IKKβ over-expression differentially regulates IVD inflammatory cytokine, chemokine, catabolic enzyme, and neurotrophic factor gene expression over time

NF-κB mediates inflammatory responses via an upregulation in inflammatory cytokine, chemokine, and catabolic enzyme expression (8, 18, 19). To assess this type of activation in IKKβCA IVDs, gene expression was assessed from RNA isolated from whole caudal IVD tissue, containing the AF, NP, and EPs. To assess for early activation of NF-κB, IVD tissue was harvested at 1-week post recombination, while sustained activation of NF-κB was assessed in IVD tissues harvested 2-months post recombination. At 1-week post recombination, gene expression of the inflammatory mediators, *Il1b* (p=0.0024)*, Il6* (p=0.023), and *Nos2* (p=0.015) were increased in IKKβCA IVDs compared to control (Fig. 3). Significant increases in catabolic enzymes, *Mmp3* (p=0.021) and *Mmp9* (p=0.028), and pro-inflammatory chemokines, *Mcp1* (p=0.00018) and *Mif* (p=0.013), were also observed in IKKβCA mice compared to control 1-week post recombination (Fig. 3). Extending this analysis of gene expression changes to 2-months post recombination, *Nos2* remained significantly increased in IKKβCA IVDs compared to control (Fig. 3). Further, at 2-months post recombination the catabolic enzymes, *Adamts4* (p=0.014) and *Mmp3* (p=0.041), and pro-inflammatory chemokines, *Mcp1* (p=0.0092) and *Mif* (p=0.011), were upregulated in IKKβCA mice compared to control (Fig. 3). In addition neurotrophic factors implicated in pathological nerve ingrowth in DD, *Ngf* (p=0.033) and *Bdnf* (p=0.040), were upregulated in IKKβCA IVDs compared to control at 2-months post recombination (Fig. 3). Together gene expression results suggest NF-κB over-activation contributed early and lasting pro-degenerative molecular changes which included an upregulation of pro-inflammatory markers, chemokines and immune cell activation mediators, catabolic enzymes, and neurotrophic factors within IVD cells.

**Figure 3:**
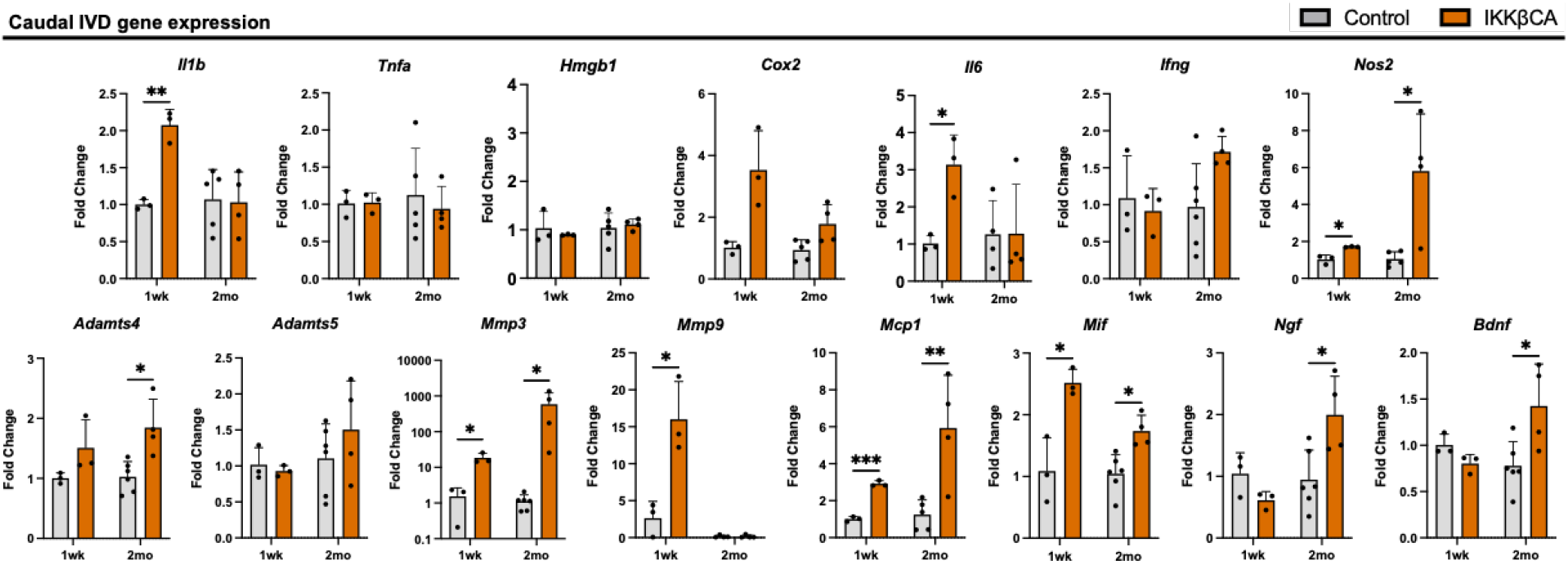
IKKβ over-expression upregulates inflammatory cytokine, chemokine, catabolic enzyme, and neurotrophic factor gene expression. Gene expression changes (relative to control) from total RNA isolated from control and IKKβCA whole IVDs containing NP, AF, and EP, 1-week and 2-months post recombination. *p<0.05, **p<0.01, ***p<0.001.

### NF-κB over-activation in IVD cells produces severe morphological IVD degeneration and disrupts functional mechanical properties

The IVD is a composite fibro-cartilaginous connective tissue structure consisting of distinct tissue types: the AF is a highly fibrous outer ring which encompasses a gelatinous proteoglycan rich NP, and the cartilage endplates (EPs) anchor the IVD to the adjacent vertebrae. To determine the effect of NF-κB mediated inflammation on IVD integrity, caudal IVD motion segments were assessed for degenerative histomorphological changes (24). At 1- and 2-months post recombination, mild degenerative changes within the NP (reduced cellularity and increased cell clustering) and AF (loss in concentric lamellae structure and cellularity) could be seen in IKKβCA mice but not control IVDs (Fig. 4A). However, these mild observable differences did not produce statistically significant changes in histological grading scores between groups at 1- or 2-months (Fig. 4A). Interestingly, IKKβCA mice at 4- and 6-months post recombination showed histological characteristics of severe DD. Compared to control mice, IKKβCA mice showed decreased cellularity, loss of Safranin-O staining, loss of NP tissue structure and disruption of the border that differentiates the NP from the AF regions (i.e. NP-AF border) (Fig. 4A). AF tissue integrity was also altered, including the presence of rounded cells near the NP-AF border and inner half of the AF, loss of cellularity in the inner half of AF, and loss of lamellae structure or widened lamellae (Fig. 4A).

**Figure 4:**
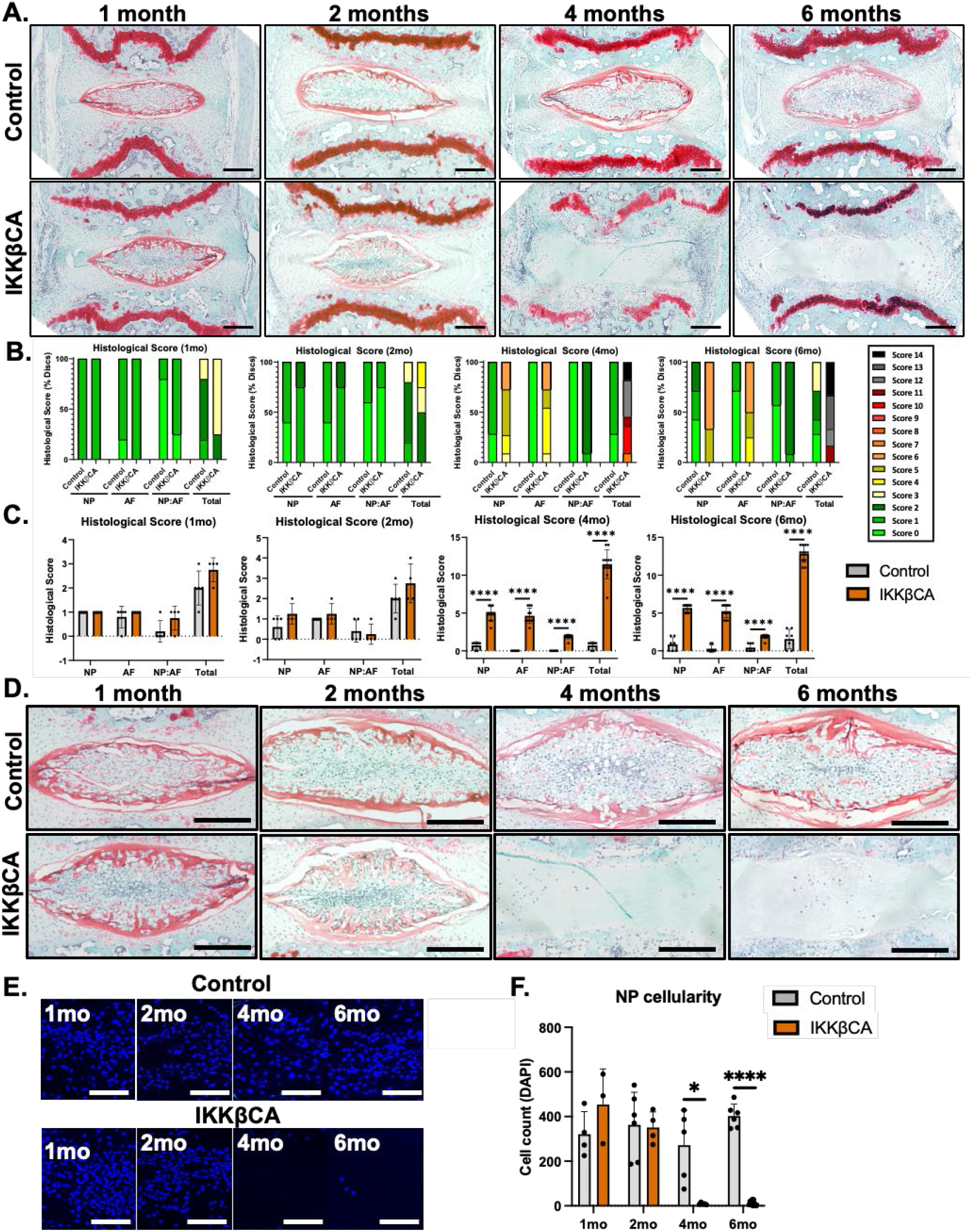
IKKβ over-expression produces severe DD. (**A**) Representative images of safranin-O stained mid sagittal sections of control and IKKβCA IVDs 1-, 2-, 4-, and 6-months post recombination. Scale bar = 250μm. Histological scoring legend, ranging from 0 (healthy) to 14 (most severe). (**B**) Distribution of histological scores. (**C**) Histological scoring within NP, AF, and NP:AF border compartments, and total score. (**D**) Representative images of safranin-O stained mid sagittal sections of control and IKKβCA IVDs at 1-, 2-, 4-, and 6-months post recombination. Scale bar = 250μm. (**E**) Representative images of DAPI (nuclear) stained mid sagittal sections. Scale bar = 100μm. (**F**) Quantification of NP cellularity within hand drawn ROIs of DAPI nuclear stained mid sagittal sections. *p<0.05, ****p<0.0001.

Degenerative changes were not detected in IVDs of control mice. This led to significant increases in histological scores in IKKβCA mice across all scoring criteria at 4-(p<0.0001) and 6-months (p<0.0001) post recombination compared to control (Fig. 4B,C).

Among the defined histomorphological characteristics of degenerative IVD, a striking change following prolonged NF-κB pathway activation was the loss of NP cellularity compared to IVDs of control mice (Fig. 4D). While no significant changes in NP cell count were detected between IKKβCA and control mice at 1- and 2-months, significant decrease in NP cell count was detected in IKKβCA mice at 4- (p=0.018) and 6-months post recombination (p<0.0001) compared to control mice (Fig. 4E,F).

Degenerative EP changes in cellularity, hyaline cartilage organization, clefts and micro fractures, bony sclerosis and tissue defects were also evaluated based on an EP specific grading criteria (25) that was not part of the IVD histologic scoring criteria used for evaluating the NP and AF. Degeneration within the EP were observed in IKKβCA IVDs at 4- and 6-months post recombination, including clear cartilage and cellular disorganization (Fig. 4A). Furthermore, potential infiltrating cell populations were detected within the EP of IKKβCA IVDs at 4- and 6- months, as indicated by nuclear stain, which was not observed in control IVDs (Fig. 4A). Lastly, degeneration within the adjacent GPs of IKKβCA IVDs was also observed at 4- and 6-months and included GP erosion and discontinuity (Fig. 4A). These results suggest that all tissue compartments of the IVD and the closely associated GP are severely affected by prolonged NF-κB activation.

To determine if the observed degeneration was associated with ECM changes, specific histological staining was used to evaluate changes in GAGs (Alcian blue) and collagen (Picrosirius red) content. Compared to control mice, a mild decrease in GAG staining intensity was observed in the inner AF of IKKβCA mice at 2-months, while a more notable loss was observed throughout the IVD of IKKβCA mice at 4-months post recombination (Fig. 5A). However, at 6-months post recombination IKKβCA mice exhibited an NP that lacked the notochordal cell pattern and was replaced by an acellular disorganized matrix that stained positively for GAGs (Fig 5A). In the AF, an increase in GAG staining in the pericellular matrix of large, rounded AF cells was observed in 6-month IKKβCA IVDs (Fig. 5A). Evaluation of collagen staining revealed no major differences between IKKβCA and control IVDs at 2-months post recombination (Fig. 5B). Whereas at 4- and 6-months post recombination, more pronounced collagen staining was detected within the NP and irregular lamellar AF structures of IKKβCA IVDs compared to control IVDs (Fig. 5B). The differences in GAG and collagen staining suggest prolonged activation of NF-κB contributes to ECM remodeling and displacement of AF tissue into the NP region, which transitioned into a fibrotic-like NP structure. Overall, histological assessment demonstrated that IVD IKKβ-NF-κB over-activation leads to severe DD, consistent with features of human DD.

**Figure 5:**
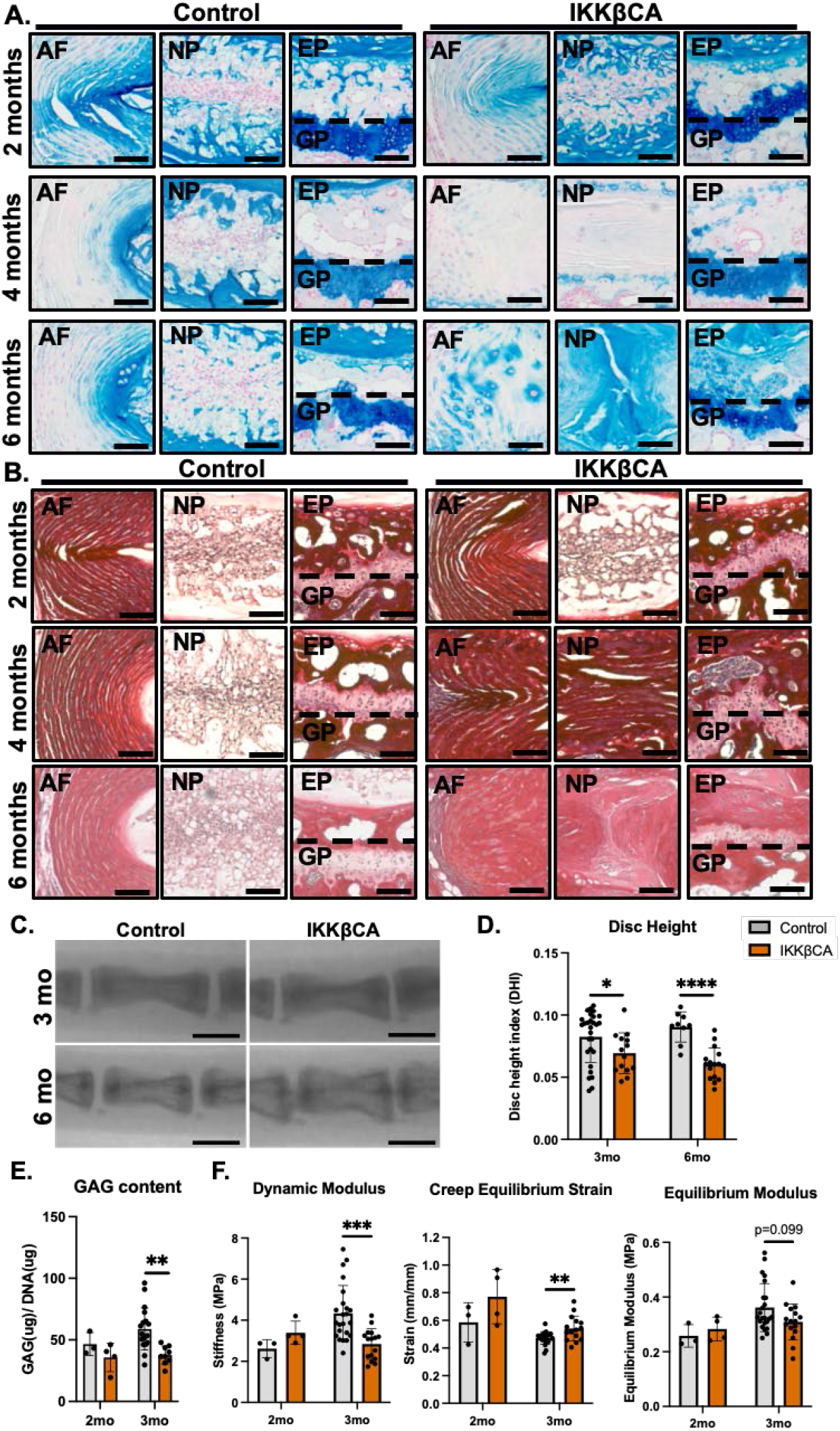
IKKβ over-expression mediates a loss of ECM, disc height, and weakened mechanical properties. (**A**) Representative Alcian blue (GAG) and (**B**) Picrosirius red (collagen) stained images of control and IKKβCA IVDs mid sagittal sections at 2-, 4-, and 6-months post recombination. Scale bar = 100μm. (**C**) Representative fluoroscopic images of control and IKKβCA C6-C8 IVDs 3- and 6-months post recombination. Scale bar = 1mm. (**D**) IVD height quantified via DHI of control and IKKβCA discs 3- and 6-months post recombination. (**E**) GAG content (μg) normalized to total DNA content (μg) within control and IKKβCA IVD digests 2- and 3-months post recombination. (**F**) Dynamic modulus (MPa), creep equilibrium strain (mm/mm) and equilibrium modulus (MPa) of control and IKKβCA discs 2- and 3-months post recombination. *p<0.05, **p<0.01, ***p<0.001, ****p<0.0001.

Having observed structural and compositional changes, changes in ECM biochemistry, functional biomechanics, and IVD height with prolonged inflammation were evaluated at time points prior to (2-months post recombination) and during the onset of morphological degeneration (3-months post recombination). For this, whole IVDs were digested for a quantification of GAG content (normalized to total DNA content). The GAG content was found to be similar in IKKβCA and control IVDs (Fig. 5E) at 2-months, consistent with the mild degenerative structural changes observed. However, at the 3-month time point, a significant loss in GAG content was observed in IKKβCA IVDs compared to control (p=0.0033, Fig. 5E). Quantification of GAG loss within IKKβCA IVDs is consistent with differences in GAG staining, and further demonstrates that localized NF-κB over-activation mediates ECM degradation within the IVD.

Furthering the functional evaluation, unconfined compression testing was performed using two impermeable platens, where dynamic modulus (MPa), equilibrium strain (mm/mm) and equilibrium modulus (MPa) were calculated during 20 cycles of dynamic loading to 0.25N (1x body weight) followed by an applied static load of 0.25N. The dynamic modulus of IKKβCA IVDs at 3-months post recombination was lower than that of control (p=0.00058, Fig. 5F). Creep testing revealed an increase in creep equilibrium strain in 3-months post recombination IKKβCA IVDs compared to control (p=0.0097, Fig. 5F). This was associated with a trend change in the equilibrium modulus within IKKβCA IVDs compared to control (p=0.099) (Fig. 5F). Prior to structural changes, at 2-months post recombination, mechanical testing revealed no significant differences in dynamic modulus, creep equilibrium strain, or equilibrium modulus between groups (Fig. 5F). Ultimately, changes in compressive properties of IKKβCA IVDs revealed a loss of dynamic compression functionality and resistance to compressive loading.

In another functional output, fluoroscopy imaging was utilized to assess changes in the disc height index (DHI), commonly used as a clinical indicator of DD (26). DHI analysis of caudal spines from IKKβCA and control mice showed discernable qualitative decreases in IVD height between caudal vertebrae of IKKβCA mice compared to control mice (Fig. 5C). Quantitative analysis revealed a significant decrease in caudal DHI in IKKβCA mice compared to control mice at 3- (p=0.044) and 6-months (p<0.0001) post recombination (Fig. 5D). The loss of IVD height in IKKβCA caudal spine reveals another functional consequence of prolonged NF-κB activation possibly associated with the loss of GAG content.

### NF-κB over-activation in IVD cells promotes macrophage recruitment and activation

In addition to IVD tissue degeneration, H&E staining revealed increased cellularity in the outer AF regions of IKKβCA mice at all time points (Fig. 4A, S2). In quantitative analysis of cell nuclei in the outer AF regions, compared to control mice, IVDs from IKKβCA mice had significantly increased cellularity at 1- (p=0.028), 2- (p=0.00020), 4- (p=0.028), and 6-months (p=0.046) post recombination (Fig. S2). To further investigate the identity of these cell populations, we performed immunostaining for the pan-macrophage marker F4/80. Additional phenotyping of these cells using CD38 as a marker of pro-inflammatory (M1) macrophages and CD206 as a marker of the inflammatory-resolving (M2) macrophage were performed. Increased expression level of F4/80*^+^* cells was observed within IKKβCA AF compartments across all time points (1-, 4-, and 6-months post recombination, p<0.0001), and within IKKβCA EP compartments at 4- and 6-months (p<0.0001) post recombination when compared to control IVDs (Fig. 6A,B). Cells expressing the M1 marker, CD38, increased within IKKβCA AF compartments at 1- (p=0.0013), 4- (p<0.0001), and 6-months (p=0.0029) post recombination, and within IKKβCA EP compartments at 4- (p<0.0001) and 6-months (p<0.0001) post recombination when compared to control IVDs (Fig. 6A,B).

**Figure 6:**
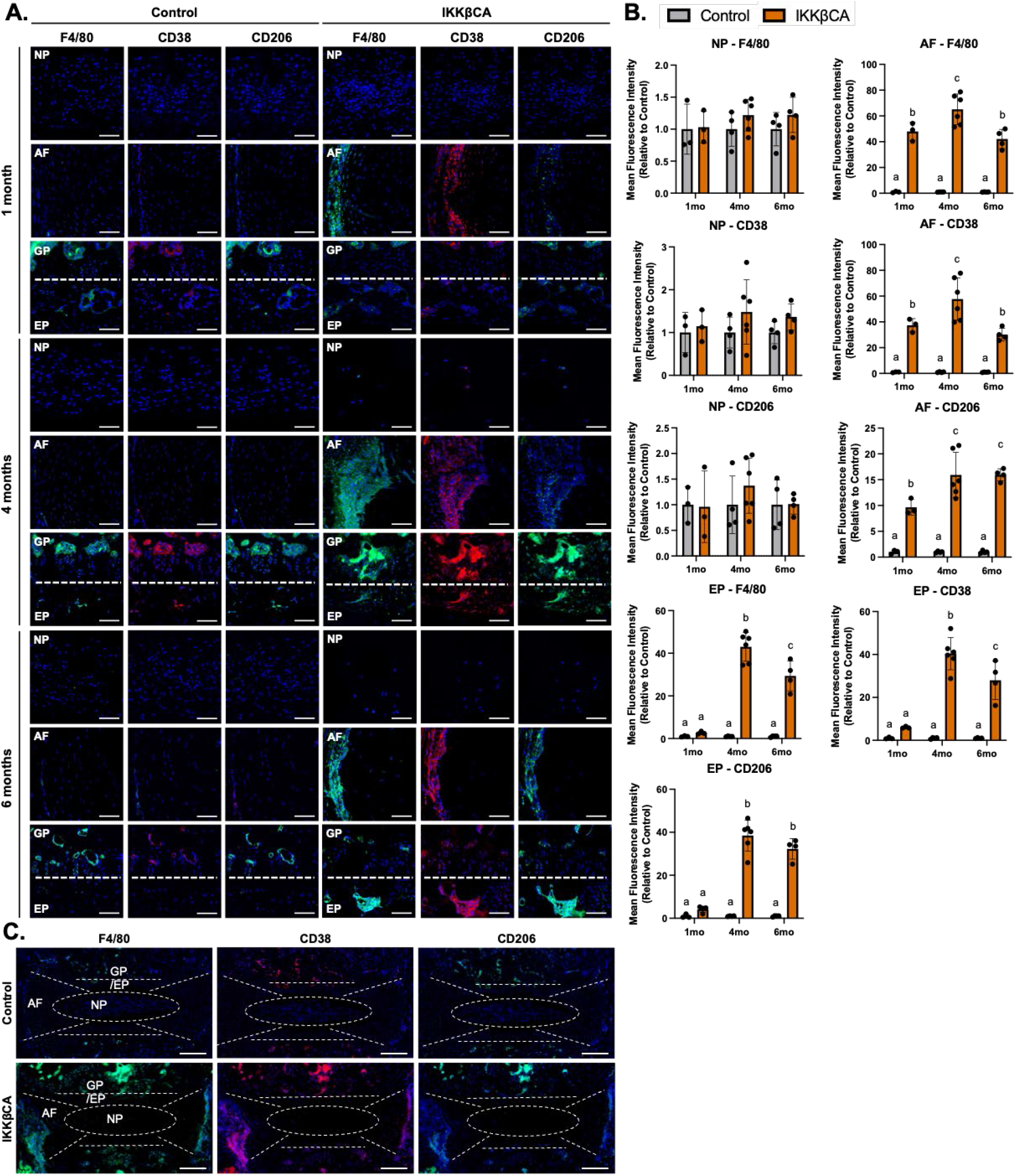
IKKβ over-expression increases macrophage presence within the IVD. (**A**) Representative images of IF staining for F4/80, CD38, and CD206 in mid sagittal sections of control and IKKβCA discs 1-, 4-, and 6-months post recombination. Scale bar = 100μm. (**B**) MFI quantification of F4/80, CD38, and CD206 expression within individual NP, AF, and EP compartments. Letters (a,b,c) indicating statistically significant (p<0.05) different groupings. (**C**) Representative images of whole disc sagittal sections of control and IKKβCA discs 4-months post recombination with delineation of tissue compartments. Scale bar = 200μm.

Expression of both F4/80^+^ and CD38^+^ cells peaked within AF and EP compartments of IKKβCA IVDs at 4-months before slightly but significantly decreasing at 6-months post recombination (Fig. 6A,B). Expression of CD206^+^ cells also significantly increased in IKKβCA IVDs over time, however the temporal pattern of expression of CD206^+^ cells differed from CD38^+^ cells. CD206^+^ cells significantly increased in IKKβCA AF compartments at 1- (p=0.0039), 4- (p<0.0001), 6- months (p<0.0001) post recombination and within IKKβCA EP compartments at 4- (p<0.0001) and 6-months (p<0.0001) post recombination when compared to control IVDs (Fig. 6A,B). No significant differences in any macrophage marker expression was observed within the NP compartments between IKKβCA and control IVDs (Fig. 6A,B). These results suggest NF-*κ*B overactivation within IVD cells, and the associated molecular changes, initiate an immune response by recruiting and maintaining a population of macrophages within the AF and EP regions of the IVD. The IKKβCA microenvironment promoted a greater abundance of pro-inflammatory (CD38^+^) over inflammatory-resolving (CD206^+^) macrophage subsets in the IVD, based on immunofluorescence imaging, suggesting that the IKKβCA IVD promoted a more pro-inflammatory microenvironment.

### The pro-inflammatory effects of IVD cell secretome are mitigated by the secretome of inflammatory-resolving macrophages

To identify possible mechanisms responsible for the recruitment and polarization of inflammatory macrophages, we analyzed the secretome released by IKKβCA IVD cells following recombination using whole IVD organ culture (Fig. 7A). Conditioned media (CM) from IKKβCA IVD organ cultures had significantly greater levels of inflammatory cytokines, IL-1β, IL-6, and IFN-γ and the chemokine, MCP-1, compared to CM from control IVDs (Fig. 7A,B).

**Figure 7:**
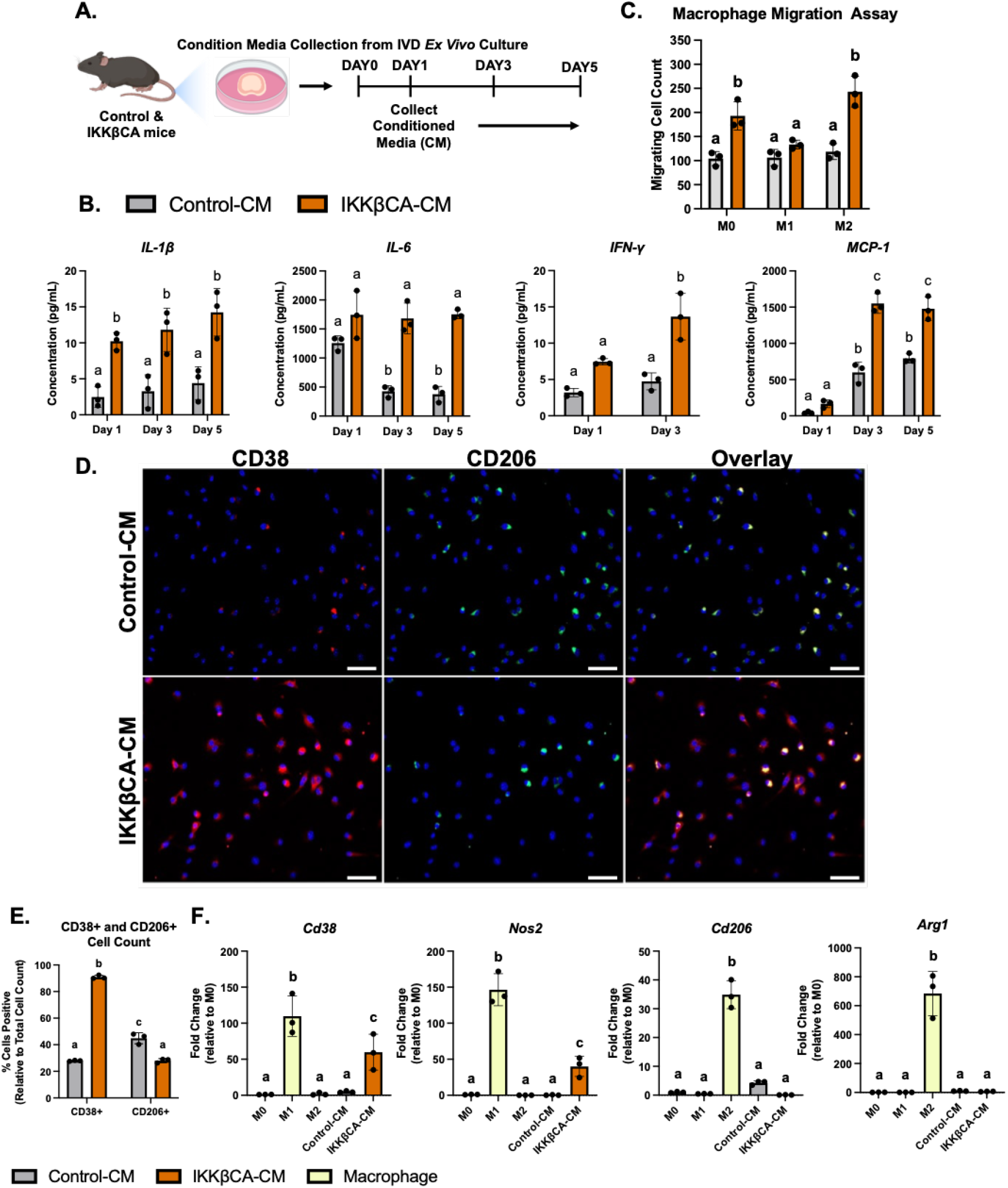
IKKβCA-CM increases macrophage migration and polarization towards an inflammatory phenotype. (**A**) Study design schematic of whole organ *in vitro* culture and conditioned media collection. (**B**) Protein concentrations (pg/mL) within conditioned media analyzed after 1-, 3-, and 5-days in culture. Letters (a,b,c) indicating statistically significant (p<0.05) different groupings. (**C**) Quantification of M0, M1, and M2 macrophage migration through a transwell membrane via DAPI nuclear count. (**D**) Representative images of IF staining for CD38 and CD206 within BMDMs cultured in 2D monolayer and stimulated with control or IKKβCA conditioned media. Scale bar = 100μm. (**E**) Quantification of % positivity for CD38 and CD206 within BMDMs treated with conditioned media. (**F**) Gene expression of M1 and M2 phenotypic markers within M0, M1, or M2 macrophages in basal media, or M0 macrophages with or without IVD conditioned media stimulation. Letters (a,b,c) indicating statistically significant (p<0.05) different groupings.

Next, in evaluating functional and phenotypic changes of macrophages exposed to IKKβCA IVD, we investigated macrophage migration behavior toward control or IKKβCA-CM. Macrophage migration across a transwell membrane (8μm) was quantified following culture with each CM. We saw an increase in naïve (M0) (p=0.0028) and M2 (p=0.0001) cell migration to IKKβCA-CM when compared to control-CM (Fig. 7C, Fig. S3). No difference was observed in M1 macrophage migration (p=0.64) (Fig. 7C). To further investigate the effect of IKKβCA microenvironment on macrophage phenotype, phenotypic markers for M1 (CD38, NOS2) and M2 macrophages (CD206, ARG1) were evaluated following *in vitro* stimulation of naïve macrophages with IKKβCA or control IVD-CM. In IKKβCA-CM, cells positive for CD38 were significantly higher (p<0.0001) while cells positive for CD206 (p<0.0001) were significantly lower compared to control-CM conditions (Fig. 7D,E). At the gene expression level, M1 macrophage markers *Cd38* (p=0.0123) and *Nos2* (p=0.0117) were significantly increased in M0 macrophages stimulated with IKKβCA-CM compared to control-CM, while no significant changes were observed for the M2 macrophage markers *Cd206* and *Arg1* (Fig. 7F). *In vitro* results support the *in vivo* findings where NF-κB over-activation initiated inflammatory molecular changes in IVD cells which directly increased macrophage migration and polarization towards an M1 phenotype while suppressing M2 polarization, possibly through secreted inflammatory mediators, IL-1β, IL-6, IFN-γ.

To identify whether the inflammatory macrophage phenotype induced by IKKβCA IVDs could be reversed, we evaluated the effect of subsequent treatment with an inflammatory-resolving M2 secretome on macrophages initially stimulated by the IKKβCA-CM or control-CM (Fig. 8A). Macrophages treated with basal media following IKKβCA-CM stimulation exhibited the same level of positivity for CD38 and CD206 as macrophages maintained in IKKβCA-CM for the duration of the experiment. This suggests that washout of IKKβCA-CM is not sufficient to eliminate or reverse the pro-inflammatory phenotypic state (Fig. 8B,C). However, when cells from IKKβCA-CM group are subsequently treated with M2-CM, a significant decrease in CD38+ cells and significant increase in CD206+ cells was observed (p<0.0001) (Fig. 8B,C).

**Figure 8:**
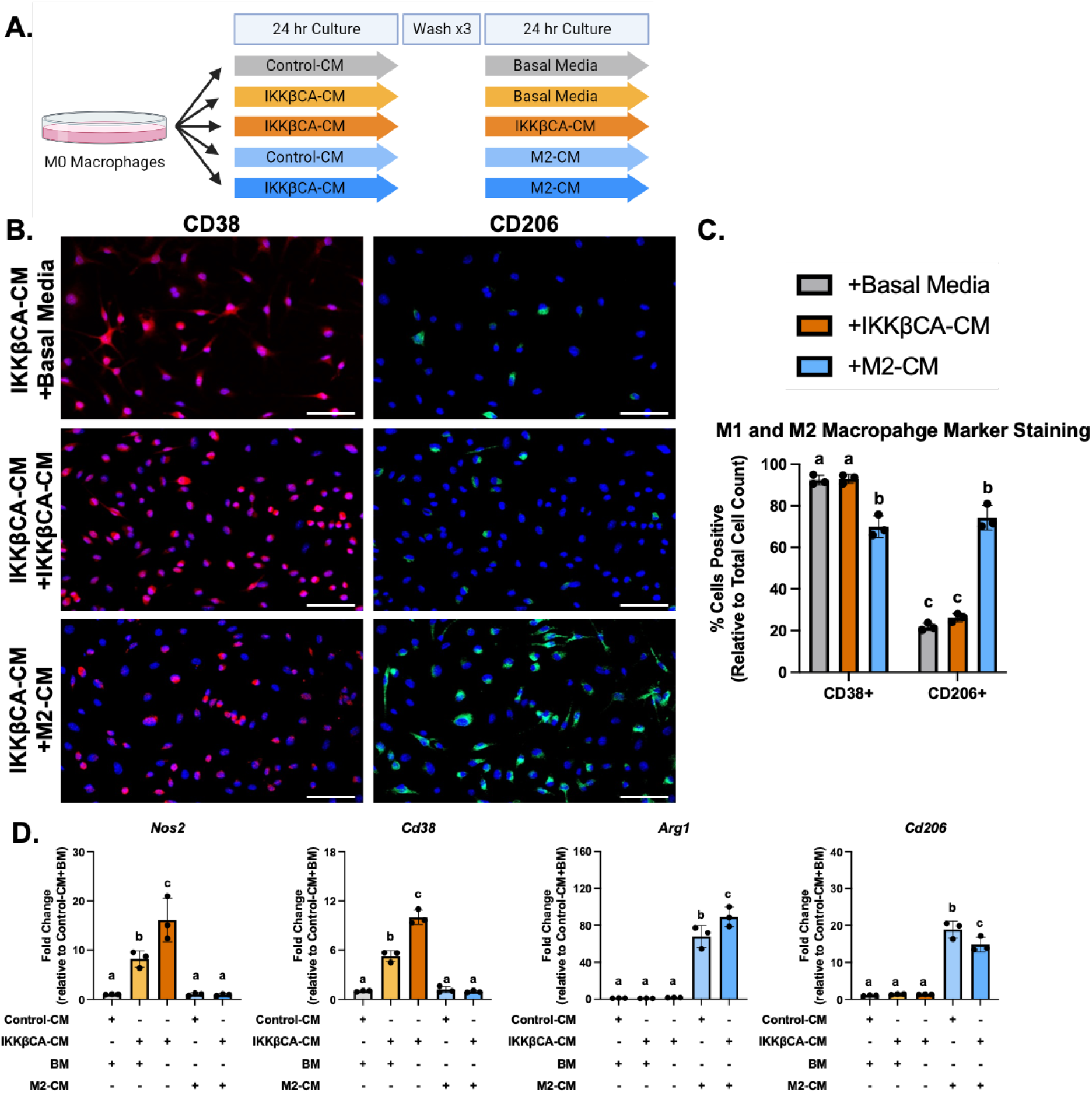
IKKβCA-CM mediated macrophage inflammatory responses are mitigated by co-stimulation with an M2 secretome. (**A**) Study design schematic for CM stimulation of M0 macrophages. (**B**) Representative images of IF staining for *CD38* and *CD206* within BMDMs cultured in 2D monolayer and stimulated with IKKβCA-CM followed by basal media or M2 macrophage CM. Scale bar = 100μm. (**C**) Quantification of % positivity for *CD38* and *CD206* within BMDMs following CM stimulation. (**D**) Gene expression of M1 and M2 phenotypic markers within M0 macrophages following CM stimulation. Letters (a,b,c) indicating statistically significant (p<0.05) different groupings.

Gene expression analysis results further support the findings, where M2-CM significantly decreased expression of *Nos2* (p<0.0001) and *Cd38* (p<0.0001), and increased expression of *Arg1* (p<0.0001) and *Cd206* (p<0.0001), when compared to groups treated with IKKβCA-CM for duration of the study (Fig. 8D). Basal media treatment following IKKβCA-CM also decreased *Nos2* (p<0.01) and *Cd38* (p<0.0001) expression compared to IKKβCA-CM treatment for duration of the study, though expression levels remained significantly higher than cells treated with M2-CM following IKKβCA-CM (p<0.05) (Fig. 8D). A greater increase in *Arg1* was achieved in M2-CM treatment following IKKβCA-CM compared to M2-CM treatment of control cells (p<0.0001) (Fig. 8D). Ultimately, these results suggest that M2-CM can reverse inflammatory polarization of macrophages activated by IVD cells over-expressing IKKβ, highlighting a therapeutic potential of M2 macrophages in initiating this resolution.

## DISCUSSION

Utilizing an *AcanCreERT2;Ikkβca* mouse model, our findings indicate that prolonged NF-κB activation in IVD cells leads to severe structural degeneration with a complete loss of NP cellularity, loss of GAG content, shift of notochordal NP to a cartilaginous NP, and eventual fibrosis of the NP. These changes were associated with functional loss of IVD height and compressive mechanical properties. IVD structural changes were accompanied by increased macrophage recruitment and increases in gene expression of inflammatory cytokines, chemokines, catabolic enzymes, and neurotrophic factors within IVD tissue. These findings fill in a gap in evidence on the effects of prolonged inflammatory activation on IVD integrity, and support the utility of the *AcanCreERT2;Ikkβca* mouse to address key questions and evaluate therapeutics for treatment of persistent IVD inflammation and degeneration.

While macrophages are the most prominent immune cell present in degenerated human discs (27–29), there are conflicting reports about macrophage phenotype found in human DD. One study found both M1 (CCR7+) and remodeling M2c (CD163+) cells increased with human DD severity, particularly in regions with structural irregularities and defects (28). However, other human analyses found M2 cells to decrease significantly with degeneration while the proportion of M1 polarized cells increased (30). In examining mechanisms contributing to severe DD within IKKβCA IVDs, we observed increased presence of macrophages (F4/80+) within the outer AF of IKKβCA mice, which consisted of M1 and M2 cell subsets, with an overall higher prevalence of M1 (CD38+) over M2 (CD206+) cells. Interestingly, the ratio of M1-to-M2 cells in IKKβCA IVDs varied over time, with the highest levels occurring at 1- and 4-months post activation. However, by 6-months, the levels of M1 cells decreased significantly from peak levels, while the M2 cell levels remained elevated, resulting in an overall decrease in the M1-to-M2 ratio. These findings provide longitudinal evidence for the existence of temporal regulation of macrophage polarization in DD *in vivo*. Since signals encountered within the microenvironment drive the polarization of macrophages into distinct subsets (31), the findings on temporal changes in macrophage subsets suggests that macrophages may play a role in resolving the inflammatory cascade and promoting fibrosis in the IVD, which was indeed present in IKKβCA IVDs. Interestingly, the shift toward increased pro-resolution occurred after the onset of substantial NP cell loss and matrix degradation, however the microenvironment was not devoid of M1 cells. Our results of dynamic macrophage populations during DD are consistent with prior studies identifying the presence of both M1 and M2 cells in a degenerative IVD (15). A puncture injury model of murine DD, where early recruitment of M1 macrophages followed by a delayed but sustained recruitment of M2 macrophages was observed using flow cytometry (16). The results of the current study, therefore extend findings in the literature regarding macrophage phenotype dynamics and localization across different tissue compartments in IVDs undergoing degeneration, in a more clinically relevant model than puncture wounding of the IVD. Whether this is the result of the heterogeneity of the macrophage population driven by phenotypic shift vs recruitment of varying cell subsets remains unknown.

Secretome analysis from IKKβCA IVDs provided evidence that soluble mediators are participating in the recruitment and activation of macrophages. We observed an increase in inflammatory mediators (IL-1β, IL-6, IFN-γ) and chemokines (MCP-1), which mimicked changes observed at the gene level. The IKKβCA IVD secretome also directly activated macrophages via an increase in migration and increased inflammatory phenotypic gene (*Arg1*, *Cd38*) and cell surface markers (CD38), while decreasing anti-inflammatory phenotypic surface markers (CD206). Similar inflammatory inducing IVD-macrophage crosstalk has been observed within multiple *in vitro* studies where degenerated bovine (32) and human (33) IVD tissue was seen to polarize human macrophages towards a pro-inflammatory phenotype via increases in various inflammatory markers. Secretome analysis reveals possible chemokine targets, MCP-1 or MIF, which were found to be released by inflamed IVD cells and whose inhibition may ultimately mitigate degenerative changes mediated by infiltrating macrophages.

Coinciding with the histological degenerative changes in this model, we observed increases in *Mmp3*, *Mmp9*, and *Adamts4* gene expression, losses in GAG staining and content, and loss of IVD height within IKKβCA IVDs, which demonstrate a highly catabolic and degenerative tissue microenvironment. In a possible mechanistic role, macrophages have been seen to exacerbate *in vitro* inflammatory driven catabolic responses of rat IVD cells through an upregulation of inflammatory cytokine (*Cox2*) and catabolic enzyme (*Mmp3*, *Adamts4*) gene expression (34). Further, in another murine model examining IL-1β mediated inflammation and IVD, global IL-1ra deficiency resulted in increases in catabolic enzyme expression and degenerative ECM changes, though the presence of infiltrating immune cells was not evaluated (21). Together, the transcriptional changes and inflammatory macrophage activation observed within this model suggest a possible macrophage mediated mechanism driving the catabolic environment leading to severe DD.

Due to their plastic nature, macrophages provide a dynamic therapeutic opportunity for harnessing the associated inflammatory-resolving and regenerative functions. This has been explored in the context of the IVD, where stimulation with human M2 macrophage CM mitigated inflammatory responses initiated by TNFα in human NP cells (35). Similarly, our results saw stimulation with an M2 secretome to reverse the inflammatory macrophage responses induced by IVD cells over-expressing IKKβ. Ultimately adding strong support for harnessing the potential of an M2 macrophage within a chronically inflamed IVD tissue.

The results presented here are consistent with prior studies utilizing NF-κB activation to study the role of persistent inflammation in musculoskeletal disease pathology, specifically tendinopathy and knee OA, and extend the understanding on immune cell crosstalk in musculoskeletal degeneration. Abraham et al. observed IKKβ mediated NF-κB over activation within tendon fibroblasts contributed to rotator cuff tendon degeneration and impaired healing driven by an upregulation in pro-inflammatory cytokines (5). Further, Catheline et al. saw IKKβ mediated activation of the NF-κB subunit, p65, to accelerate a degenerative OA phenotype within articular cartilage driven by local pro-inflammatory secretory factors (36). Inversely, studies targeting the inhibition of NF-κB activation via genetic deletion or siRNA knockdown of p65 within murine models observed a protection from OA progression (37, 38). Though not all studies evaluated immune cell infiltration, increased immune cell presence was observed within the tendon and synovium of NF-κB over-expression models, however crosstalk between the inflamed tissue and immune cells was not evaluated (5, 36). Ultimately our work provides additional support for persistent inflammation being a primary cause of musculoskeletal disease progression while also adding novel insight into the role inflamed IVD cells play in recruiting and activating macrophages.

Limitations to this study include the well-known differences between murine and human IVDs (39). Nonetheless, findings from *in vivo* animal models inform relevant biological processes that may contribute to the better understanding of human DD. A second limitation is that while a similar upregulation of inflammatory cytokines, chemokines, and catabolic enzyme gene expression was observed in lumbar IVDs following NF-κB activation (Fig. S1), minimal changes were detected in lumbar IVD health (Fig. S4). Possible explanations for regional differences within IKKβCA mice could be unique structural, mechanical properties and vascularity differences between lumbar and caudal IVDs (40, 41). Although the phenotypic differences limited the scope of this study to evaluating caudal IVDs, these results inspire future research questions. Further, the time point chosen for this study might have been insufficient to capture the changes in lumbar IVDs due to NF-κB over-activation, and thus, DD in lumbar IVDs may require longer time points post recombination.

In summary the findings of this study provide evidence that prolonged canonical IKKβ-NF-κB signaling pathway leads to accelerated DD by 4-months. Model characterization showed that IKKβ overexpression within the IVD led to a decrease in IVD height, loss of IVD structure, composition, and cellularity, and loss in compressive mechanical properties. Moreover, IKKβ overexpression led to the recruitment of an activated macrophage population to the IVD. The increased production of inflammatory cytokine, chemokine, catabolic enzyme, and neurotrophic factor genes and proteins downstream of canonical NF-κB activation is believed to mediate these degenerative changes and to directly recruit and activate innate immune cells, such as macrophages. Lastly, we identified stimulation with an M2 macrophage secretome can mitigate the IVD cell driven inflammatory changes. Together these results provide support for characterizing the NF-κB mediated chronic inflammatory environment within the IVD, and provide a model for which therapeutic targets, including downstream targets of NF-κB and the utility of an inflammatory-resolving M2 macrophage, may be investigated for their potential in mediating chronic inflammation and subsequent severe DD.

## METHODS

### Mice

Procedures involving the use of animals in this study were performed after attaining approval from the Institutional Animal Care and Use Committee at Columbia University. Conditional IKKβ ‘gain-of-function’ *R26Stop^FL^ikk2ca* mice (JAX stock no. 008242) were used to induce canonical NF-κB pathway activation (42). Homozygous *Ikk2ca^fl/fl^*mice were bred to mice heterozygous for aggrecan (*Acan*) knock-in allele carrying tamoxifen-inducible form of Cre recombinase (*Acan^CreERT2/+^*; JAX stock no. 019148) (43). Mice without CreER^T2^ recombinase were used as controls (*Acan^+/+^;Ikk2ca^fl/fl^*, Control) for comparison to CreER^T2^-positive mice (*AcanCre^ERT2/+^;Ikk2ca^fl/fl^*, IKKβCA). For initial *in vivo* histopathology analysis, Cre-mediated recombination was induced in skeletally mature (3-4 months-of-age) mice via intraperitoneal (IP) tamoxifen injections (0.1 or 0.3 mg/g of body weight dissolved in sunflower seed oil; Sigma-Aldrich, Cat. No. T5648) for 5 or 3 consecutive days, respectively. CreER^T2^ negative littermate control mice received the same tamoxifen injection. IVDs were isolated from the caudal spine (C5-C10) between 1- to 6-months post IP recombination.

To evaluate Cre activity within the IVD, heterozygous *AcanCre^ERT2/+^*mice were crossed with conditional *Ai14* reporter mice (JAX stock no. 007914) to generate *AcanCre;Ai14* mice (44). Recombination was induced as described above, and IVDs were isolated from the caudal spine (C5-C10) either 3-days or 3-months following IP injection (both 0.3mg/g and 0.1mg/g of body weight doses) for assessment of localized Cre activity (Fig. 1).

### Gene Expression

Whole IVD tissues containing were isolated, snap frozen, and homogenized using a bead tissue homogenizer (Mikro-Dismembrator U, Sartorius) (n=4-6 per group). Total RNA was extracted using TRIzol and chloroform phase separation followed by RNA cleanup using spin columns (Qiagen) according to the manufacturer’s protocol. Relative gene expression was quantified normalized to glyceraldehyde-3-phosphate dehydrogenase (*Gapdh*) using the ΔΔC_T_ method (gene abbreviations and primer sequences listed in Table S1).

### Histological analysis

Caudal bone-disc-bone spine segments were fixed (4% paraformaldehyde, 24 hr), decalcified (14% EDTA, 10 days), and either processed for paraffin-embedding or soaked in sucrose and embedded in OCT for cryo-sectioning. Tissue structure was analyzed using paraffin embedded sagittal sections (7um) stained with either Safranin-O (cartilage/mucin), Alcian Blue (glycosaminoglycans, GAGs), or Picrosirius Red (Type I and III Collagen). Stained slides were imaged using an Axio Observer (Axiocam 503 color camera, Zeiss). Histomorphological analysis was performed using a previously described mouse specific histological grading system (24), with higher scores indicating increased tissue degeneration (n=4-17 samples per genotype and time point). Stained sections were scored blinded to experimental groups. Differences in histological scores between genotype groups was performed by comparing scores of IVDs collected from multiple levels in each animal.

### Cellularity measurements

Using the ImageJ software (NIH), hematoxylin (nuclear) or DAPI stained histological images were converted to 8-bit, auto-thresholded, converted to binary, and the cell number was quantified using analyze particles function within a custom-defined region of interests (ROIs) containing the outer AF or NP (45). Cell number measurements and ROI delineation were performed blinded to experimental group and analyzed comparing multi-level pooled IVDs between groups. (n=3-17 samples per genotype and time point).

### Fluoroscopy analysis

IVD height was determined for analysis of digital fluoroscopy images (Glenbrook Technologies) taken following euthanasia and prior to IVD isolation. The DHI was calculated by averaging the IVD height and normalizing to adjacent vertebral body length (n=9-28 per group and time point) (46). Measurements were taken blinded to experimental group. DHI values were analyzed comparing multi-level pooled IVDs between groups.

### Immunofluorescence microscopy and image analysis

Paraffin embedded tissue sections were baked (60°C, 35 min), deparaffinized with xylene, and rehydrated using a graded series of ethanol washes. Antigen retrieval was performed with 0.1% Triton-X (10 min). Tissue sections were blocked (45 min) for non-specific binding using background buster (Innovex Biosciences). Sections were then incubated overnight at 4°C with primary antibodies. The next day, sections were incubated for 1 hr with secondary fluorescent antibodies. Primary and secondary antibodies and dilutions are listed in Table S2. Sections were mounted with DAPI anti-fade mounting medium (Vector, H-1200) before imaging with Axio Observer (Axiocam 702 mono camera, Zeiss). Exposure settings were fixed across all tissue sections during imaging.

For protein expression quantification, fluorescence images were converted to 8-bit and auto-thresholded, and a mean fluorescence intensity (MFI) was calculated using the measurement of mean grey value function within ImageJ (NIH) software. MFI measurements were taken within custom defined ROIs (NP, AF, and EP) drawn blind to experimental groups. For nuclear MFI measurements, nuclear ROIs were created by converting DAPI stained sections to nuclear masks and measuring MFI within (n=4-12 samples per group and time point).

Cryo-embedded tissues were sectioned (7 μm) and signal recovery of the fluorescent reporter tdTomato was carried out. Sections were re-hydrated, antigen retrieval and peptide blocking was performed as detailed above. To enhance the tdTomato signal, cryo-sectioned tissues were incubated overnight at 4°C with primary antibodies followed by a 1 hr incubation with secondary fluorescent antibodies at room temperature (Table S2). Sections were mounted (DAPI) and imaged as described above (n= 3 samples per genotype and time point).

### Immunohistochemistry

Paraffin embedded tissue sections were deparaffinized as described prior. Antigen retrieval was performed using hyaluronidase solution (Sigma H3506, 100 μg/mL) at 37°C for 12 min. Endogenous peroxidase and protein blocking steps were performed using reagents provided in the ABC detection kit (Abcam, ab64261). Sections were incubated overnight with primary antibodies at 4°C (Table S2). The next day, sections were incubated with secondary antibodies and DAB staining using reagents provided in the Abcam kit, according to the manufacturer’s protocol. Sections were dehydrated, counterstained with 0.5% methyl green (Sigma, 198080), mounted with Permount mounting medium (Fisher Scientific, SP15), and imaged (Axiocam 503 color camera, Zeiss).

### Mechanical testing

IVDs were mechanically tested on a TA Electroforce DMA 3200 Mechanical Tester. Prior to mechanical testing IVDs were thawed in PBS (37°C, 1 hr). IVD height (mm) and cross-sectional area (area of an ellipse, mm^2^) of the IVDs were approximated using fluoroscopy imaging with ImageJ software. For IVD height and cross-sectional area measurements IVD samples containing the intact NP, AF, and CEPs were micro-dissected and imaged. Unconfined compression testing was performed between two impermeable platens (WinTest). A 0.02N preload was applied to each sample followed by 20 cycles of sinusoidal loading at 0.1 Hz to a maximum load of 0.25N (1x body weight). This was followed by equilibrium creep testing, where a load ramp of 0.25N was applied over 5 seconds and held for 1200 seconds. Dynamic Modulus (MPa) was calculated from the ratio of the applied stress and measured strain during the 20th cycle of dynamic loading to allow for repeatable sample displacement hysteresis. The resulting equilibrium strain (mm/mm) and equilibrium modulus (MPa: applied stress/strain) at end of the hold was measured. Measurements were analyzed using multi-level pooled IVDs between groups (n=3-18 per group and time point).

### Glycosaminoglycan Analysis

Whole IVDs were digested overnight in papain (0.3 mg/mL) in 100mM sodium acetate, 10mM cysteine HCl, and 50mM EDTA. A dimethymethylene blue (DMMB) assay was used to quantify GAG content within IVD tissue digests (47). GAG content was normalized to total DNA measured within IVD tissue digests using Pico Green Assay (n=3-18 per genotype and time point). Measurements were analyzed using multi-level pooled IVDs between groups.

### Ex vivo Culture and Generation of Conditioned Media

To isolate the effects of IVD inflammation, we established IVD *ex vivo* culture to generate CM. IKKβCA and Control mice at 1-week post recombination was sacrificed and 5 caudal IVDs were microdissected. After serial washes in phosphate buffered saline (PBS and Hank’s balanced salt solution, IVDs were cultured in DMEM/F12 with 5% Fetal Bovine Serum (FBS, Crystalgen, Cat #FBS-500HI) and 1 % Penicillin-Streptomycin. Media was changed every 2 days and CM from Day 1, 3, and 5 in culture were collected.

### Cytokine Immunoassay

Secreted cytokine levels into the collected IKKβCA- and Control-CMs were measured using 9-Plex LEGENDPlex mouse inflammation panel (BioLegend, Cat #740446) according to manufacturer’s protocol. The predefined panel enabled simultaneous quantification of 9 cytokines: CCL2 (MCP-1), GM-CSF, IFN-β, IFN-γ, IL-1α, IL-1β, IL-6, IL-10, TNF-α.

### Bone Marrow Derived Macrophage Isolation and Culture

Femur and tibia isolated from wild type C57BL/6 mice at 3 months-of-age were washed in ice cold PBS and stored in ice cold RPMI media (ThermoFisher, Cat # 11875093). Following dissection, long bones were transferred to the sterile biosafety cabinet and their bone marrows were flushed using 1mL of sterile RPMI media using 27G needle. The collected flow through was resuspended in 30 mL of complete macrophage media, containing RPMI, 10% FBS (GeminiBio, Cat #100-106), 30% L929 conditioned media (LCM), and 1 % Penicillin-Streptomycin. The cell suspension was split into three 6 cm petri dish and cultured until 80% confluency, with media change every 2 days.

### Macrophage Polarization and Transwell Migration Assay

Upon desired confluency, macrophages were chemically stimulated for 24 h to take on either M1 or M2 phenotypes. For M1 polarization, macrophages were treated with 100 ng/mL LPS (Sigma, Cat #L2630-10MG) and 20 ng/mL IFNγ (Shenandoah Biotech, Cat #200-16) in complete macrophage media. For M2 polarization, macrophages were treated with 20 ng/mL IL-4 (Shenandoah Biotech, Cat #200-18) and 20 ng/mL IL-13(Shenandoah Biotech, Cat #200-22). Cells cultured with complete macrophage media was used as M0 group.

After polarization, 1.0 x 10^5^ cells were seeded at the top of the transwell insert (8 μm pore size, Corning, Cat # 3422) positioned into the well with 500 μL of either IKKβCA-CM or Control-CM collected at Day 3 in culture. Cells were incubated in normoxia for 24 h, fixed in 4% paraformaldehyde and cells at the top of the transwell membrane were removed using cotton swab. Migrated cells located at the bottom side of the transwell membrane were permeabilized with 0.1% Triton X-100 in PBS and the transwell membrane was coverslipped with mounting medium containing DAPI (Vector Labs, Cat # H-1800-2). For each transwell, four different fields of view were imaged, and the number of migrated cells were analyzed by counting the number of DAPI-stained nuclei using ImageJ. Cell count of each transwell is an average count of four different fields of view.

### Macrophage Polarization using Conditioned Media

To investigate the effects of conditioned media on macrophage polarization, M0 macrophages were exposed to either IKKβCA-CM or Control-CM at 1:1 solution with complete macrophage media for 24 hours. Following exposure to the CMs, cells were washed with DPBS and were subjected to either RNA isolation or fixed in 4% PFA for 10 minutes for immunocytochemistry (ICC) analysis.

For sequential polarization study, M0 macrophages exposed to either IKKβCA-CM or Control-CM for 24 hours were then washed three times with DPBS and cultured in M2-CM for 24 hours. After sequential polarization, cells were harvested for RNA isolation and ICC analysis. M2-CM was generated using chemically stimulated M2 macrophages, washed three times in DPBS, cultured in complete macrophage media for 24 hours, after which the media was collected (Figure 8B).

### Immunocytochemistry

Following fixation, cells were washed with PBS, permeabilized with 0.1% Triton X-100 for 15 min. and blocked using 1% Bovine Serum Albumin (BSA) in PBS for 45 min. Samples were double stained with primary antibodies against overnight at 4°C. After incubation, cells were washed with PBS three times, and cells were incubated with secondary antibodies for 1 hr at room temperature. Following staining, cells were washed three times with PBS and mounted onto a glass slide with VectaShield DAPI mounting solution (Vector Labs, Cat # H-1800-10). For each sample, four different fields of view were imaged, and the total number of cells, and cells positive for CD38 or CD206 were manually counted. Percent cell positivity for each surface marker was calculated by dividing the number of cells positive for each surface marker by total number of cells in the field of view. Each data point of percent cell positivity is an average count of four different fields of view.

### Statistics

Differences between genotype groups were analyzed with Student’s t-test with multiple comparison correction using Holm-Šídák in Prism (V8.3.1). Differences across *in vitro* conditioned media stimulation groups were analyzed using ANOVA with multiple comparison correction using Holm-Šídák in Prism (V8.3.1). p<0.05 considered significant, and p<0.1 considered a trend.

## ACKNOWLEDGMENTS

This work was supported in part by grants from the NIH R01AR069668, R01AR077760, and R21AR080516. Flow cytometry experiments were performed in the Columbia Stem Cell Initiative Flow Cytometry core facility at Columbia University Irving Medical Center under the leadership of Michael Kissner.

## Data and materials availability

All data needed to evaluate the conclusions in the paper are present in the paper and/or the Supplementary Materials. Additional data generated and analyzed during this study may be requested from the authors.

---

**Figure S1:**
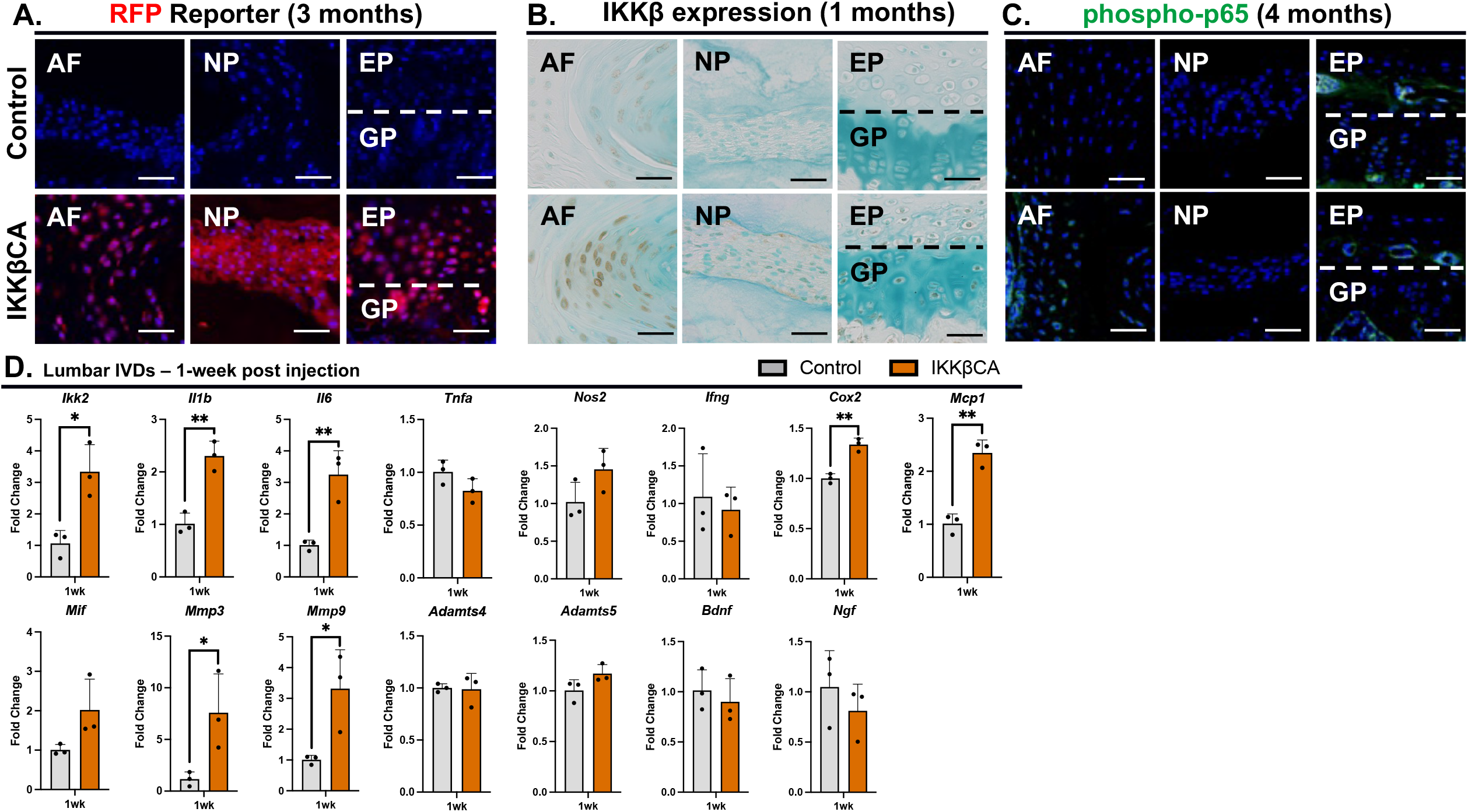
Cre-activity, NF-κB activation, and gene expression changes within lumbar IVDs. (**A**) Representative images of IF staining for RFP in control and IKKBCA mid coronal lumbar IVD sections. Scale bar = 50μm. (**B**) Representative images of IHC staining for IKKβ within mid coronal sections of 1-month control and IKKβCA lumbar IVDs. Scale bar = 50μm. (**C**) Representative images of IF staining for phospho-p65 in control and IKKBCA mid coronal lumbar IVD sections. Scale bar = 50μm. (**D**) Gene expression changes (relative to control) from total RNA isolated from control and IKKβCA whole lumbar IVDs containing NP, AF, and EP, 1-week post recombination. *p<0.05, **p<0.01.

**Figure S2:**
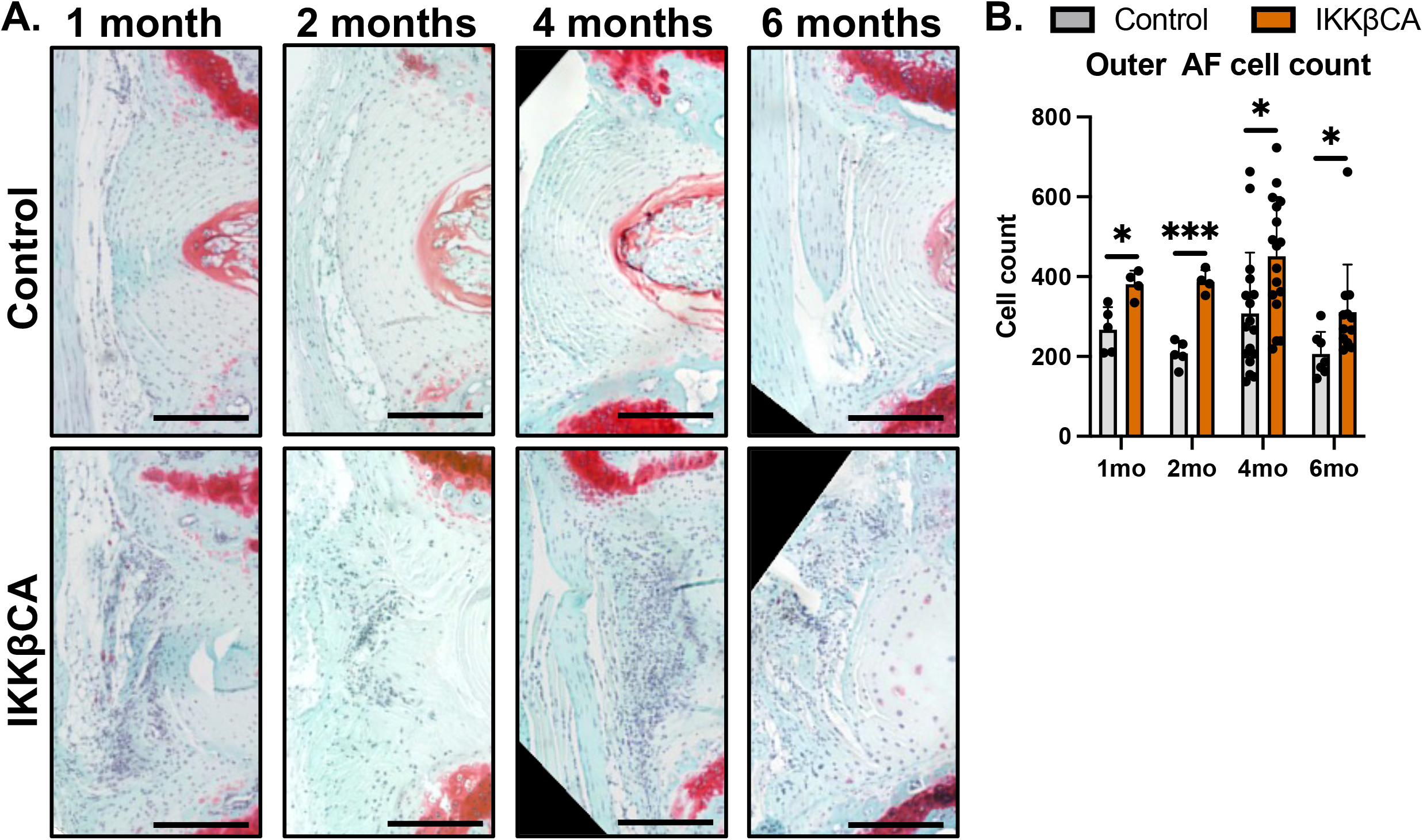
IKKβ over-expression increases cellularity presence within the AF. (**A**) Representative images of safranin-O stained mid sagittal sections of control and IKKβCA caudal IVDs 1-, 2-, 4-, and 6-months post recombination. Scale bar = 250μm. (**B**) Quantification of outer AF cellularity using haematoxylin nuclear stain.

**Figure S3:**
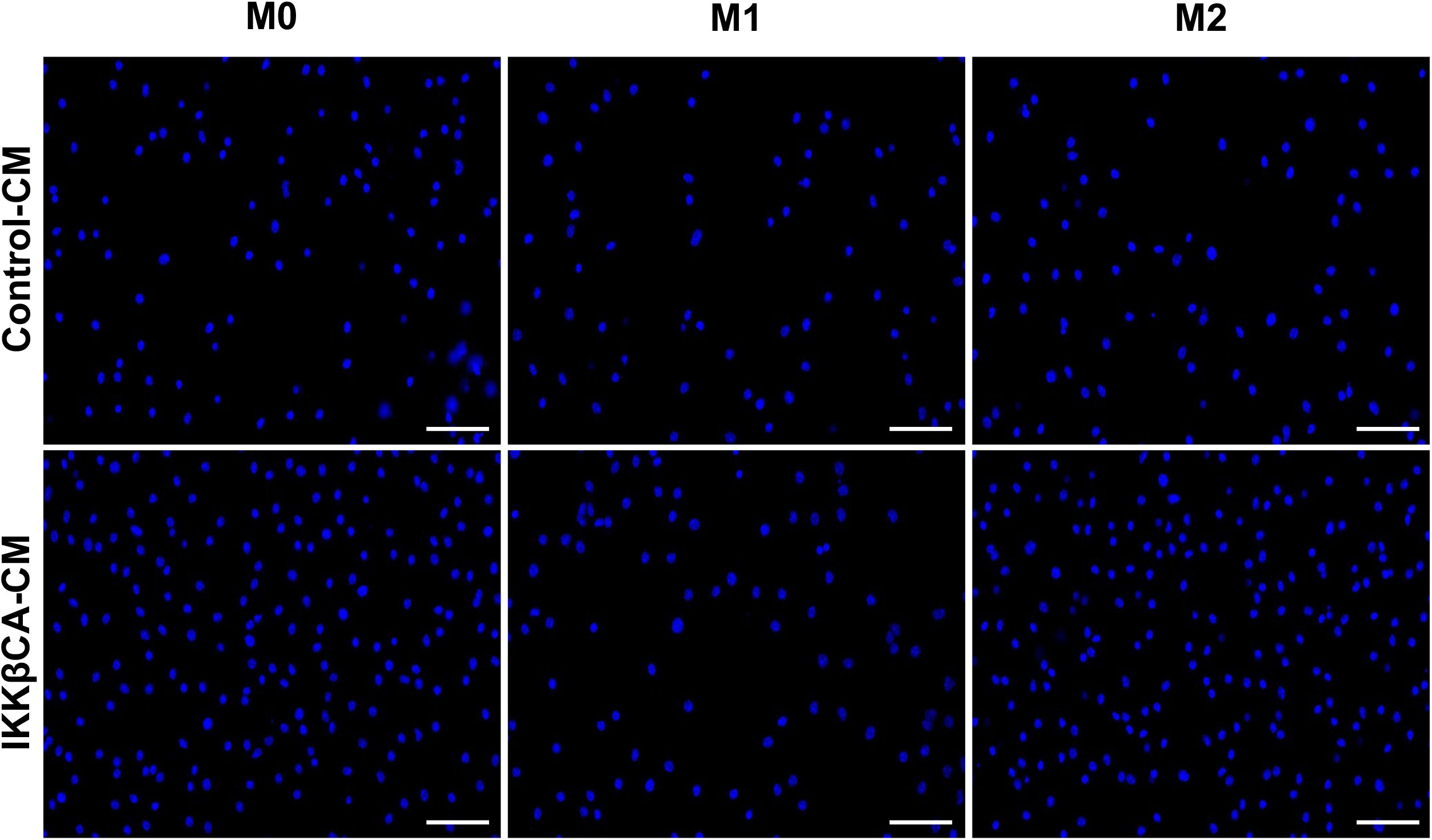
IKKβ-CM increases macrophage migration *in vitro*. Representative images of DAPI (nuclear) stained M0, M1, and M2 macrophages on transwell membranes following CM stimulation. Cell counting used for quantification of migration through transwell membranes. Scale bar = 100μm.

**Figure S4:**
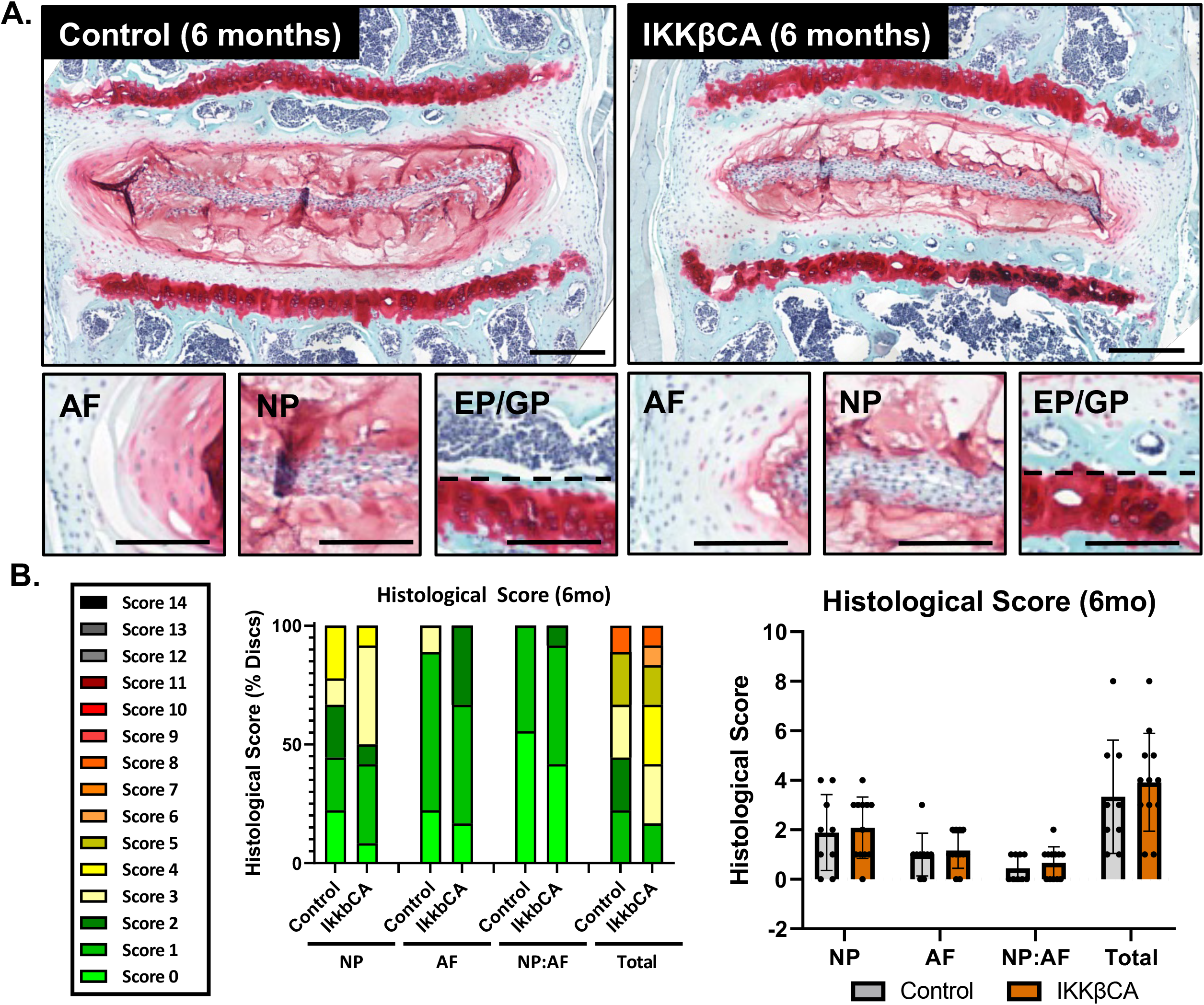
IKKβ over-expression produces no significant changes to lumbar disc health/structure at 6-months. (**A**) Representative images of safranin-O stained mid coronal sections of control and IKKBCA lumbar IVDs 6-months post recombination. Total IVD scale bar = 400μm. Compartment image scale bar = 200um. (**B**) Distribution of histological grades within NP, AF, and NP:AF border compartments, and Total score. Histological scoring severity legend ranging from 0 (healthy) to 14 (most severe). No significant differences observed.

**Table S1:**
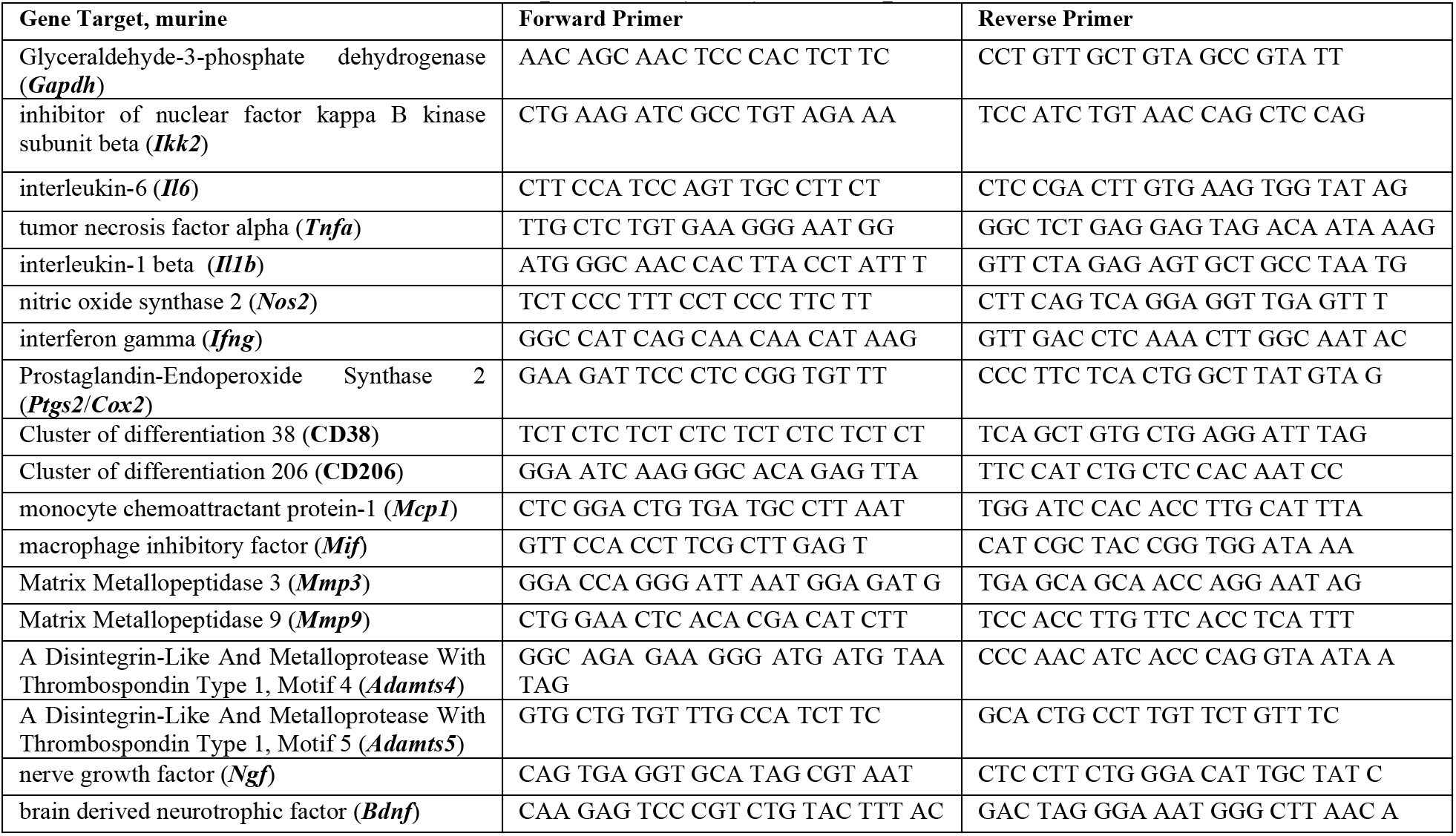
FWD and REV murine primers (IDT) for rt-qPCR.

**Table S2:** Primary and secondary fluorescent antibodies used for immunohistochemistry.

| Antigen | Company | Cat # | Dilution |
| --- | --- | --- | --- |
| [1°] mCherry | Sicgen | AB0040-200 | 1:250 |
| [1°] Phosphorylated p-65 | Abcam | AB86299 | 1:500 |
| [1°] F4/80 | Bio-rad | MCA497GA | 1:100 |
| [1°] IKKβ | Sigma | 07-1479 | 1:500 |
| [1°] CD38 | Fisher | PIMA516871 | 1:100 |
| [1°] CD206 | Fisher | PIPA595840 | 1:100 |
| [2°] Donkey anti-goat (AF 594) | Thermo Fisher | A-11058 | 1:250 |
| [2°] Donkey anti-rabbit (AF 594) | Abcam | AB150064 | 1:200 |
| [2°] Donkey anti-rat (AF 488) | Abcam | Ab150153 | 1:200 |

